# Sample size evolution in neuroimaging research: an evaluation of highly-cited studies (1990-2012) and of latest practices (2017-2018) in high-impact journals

**DOI:** 10.1101/809715

**Authors:** Denes Szucs, John PA Ioannidis

## Abstract

We evaluated 1038 of the most cited structural and functional (fMRI) magnetic resonance brain imaging papers (1161 studies) published during 1990-2012 and 273 papers (302 studies) published in top neuroimaging journals in 2017 and 2018. 96% of highly cited experimental fMRI studies had a single group of participants and these studies had median sample size of 12, highly cited clinical fMRI studies (with patient participants) had median sample size of 14.5, and clinical structural MRI studies had median sample size of 50. The sample size of highly cited experimental fMRI studies increased at a rate of 0.74 participant/year and this rate of increase was commensurate with the median sample sizes of neuroimaging studies published in top neuroimaging journals in 2017 (23 participants) and 2018 (24 participants). Only 4 of 131 papers in 2017 and 5 of 142 papers in 2018 had pre-study power calculations, most for single t-tests and correlations. Only 14% of highly cited papers reported the number of excluded participants whereas about 45% of papers in 2017 and 2018 reported excluded participants. Targeted interventions from publishers and funders could facilitate increase in sample sizes and adherence to better standards.

## Introduction

The number of participants and hence, statistical power is in general low in cognitive neuroscience and neuroimaging. Hence, false negatives, imprecise measurements, as well as exaggerated published effect sizes and high false report probability can be expected in this field (Yarkoni 2009; Ioannidis, 2008; 2005a,b; Button et al. 2013; Poldrack et al. 2017; Szucs and Ioannidis, 2017a,b).

It is often thought that statistical power is only important for studies because low power precludes the detection of existing effects. However, studies with low power also have other serious problems: First, low power increases false report probability, the probability that statistically significant findings are in fact false (Ioannidis, 2005; Szucs and Ioannidis, 2017b). Second, using low sample sizes (and therefore having low power) leads to noisy measurements due to high sampling variability. Hence, many studies with low power will likely report widely different results. Third, if mostly only null hypothesis significance testing studies (NHST) with statistically significant results are published, then these will inevitably report exaggerated (large) effect sizes even if the true phenomenon produces small effect sizes. This is so because by using small sample sizes and therefore small degrees of freedom only relatively large effects have a chance to pass traditional statistical significance testing thresholds (e.g. α = 0.05). Such large effects may occur occasionally due to sampling variability. Large effects can also be the result of p-hacking when analytical manipulation makes the results from these small studies to pass the significance threshold. Many such exaggerated published effects from small studies will then distort the literature and may also be picked up by meta-analyses, thus further resulting in exaggerated meta-analytic effect sizes.

The above makes it clear that it would be extremely beneficial for studies 1) to increase statistical power by increasing sample size and 2) to include pre-study power calculations and set sample sizes in a principled manner so that pre-defined effect sizes could be detected. Such calculations would also be required in the original null hypothesis decision framework of Neyman and Pearson (1933) so that optimal decisions with regards to rejection or no rejection of the null hypothesis could be made (for review see Szucs and Ioannidis, 2017b). In contrast to their theoretical and practical importance it is rare to see power calculations in papers published in many disciplines.

It would be useful to understand whether the problem of low power also affects the most influential neuroimaging papers and whether any improvements have occurred over time. Here, our main objective was to scrutinize participant numbers in the most cited experimental functional magnetic resonance imaging (fMRI) studies published between 1990 and 2012 and compare these to the participant numbers included in studies published in 4 top neuro-imaging journals in 2017 and 2018. Previously (Szucs and Ioannidis, 2017a) we observed that, on average, power was higher in papers in medically oriented than in cognitive neuroscience journals. So, for comparison we also report participant numbers in the most cited structural MRI (sMRI) and fMRI clinical studies that examined patients (highly cited studies were published between 1990-2012). We were especially interested in highly cited studies because (by definition) they are often cited in support of claims and because they are likely to set standards for many researchers. To monitor progress in setting participant numbers in a principled way we have also collected data about the frequency and method of (pre-study) power calculations in studies published in top neuroimaging journals in 2017 and 2018.

## Methods

### Highly cited papers: Identification and data extraction

We evaluated sample size data from 1038 of the most cited sMRI and fMRI papers (1161 studies) published during 1990-2012 and 273 papers (302 studies) published in 4 top neuroimaging journals during 2017 and 2018. By ‘paper’ we mean a publication unit published as a formal paper in a journal, whereas by ‘study’ we mean the individual studies reported in papers. Some papers reported more than one MRI study. Hence, the number of studies is higher than the number of papers. We evaluated only fMRI studies whereas some papers also included purely behavioral, electro-encephalography, and other types of non-eligible studies.

First, we queried the Scopus (scopus.com) search engine for the 1,500 most highly cited ‘articles’ using magnetic resonance imaging (MRI) published from 1990 onwards in the ‘neuroscience’ field. The date of query was 25 May 2017 and it returned papers published between 1990 and 2012. The search term was TITLE-ABS-KEY (*MRI*) AND DOCTYPE (ar) AND PUBYEAR > 1989 AND (LIMIT-TO (SUBJAREA, “NEUR”)). The query (see main text) generated a comma separated text file. During the process of data extraction we added additional records to this file describing participant numbers and study types.

We aimed to examine primary empirical research reports that used in vivo sMRI or fMRI to study brain structure and function in humans. So, we excluded misclassified review papers, methodological papers, meta-analyses of published findings, post-mortem studies, case studies, animal studies, behavioral papers, theoretical papers, modelling papers, papers on surgery which only used MRI to aid surgery, non-brain MRI papers (e.g. MRI of the chest and muscles), and papers with other than MRI technology (positron emission tomography, electro-encephalography, computed tomography).

Specifically, we first read titles and abstracts queried from the Scopus database. For all studies of interest we accessed full text pdf files where possible and we confirmed whether a certain paper was appropriate for study. If a paper was appropriate for study then we manually extracted participant numbers from most papers by reading the ‘Participants’, or equivalent, sections of full text pdf files. In case of uncertainty about participant numbers other sections of papers were also examined. We could not access pdf files for 48 relevant papers but we were able to extract participant information from abstracts and online full texts. We could not access participant data for 9 relevant papers, so they were not considered for analysis (marked ‘xnoacc’ in the data file).

In remaining sample there were 1098 papers (1223 studies). The journals most represented in our sample are shown in **Supplementary Table 1.** These studies could be sorted into 6 major categories:

(1) Experimental fMRI cognitive neuroscience studies with normal adults (experimental studies). The primary concern of these studies was the understanding of brain structure and function and they did not have primary clinical relevance. Most of the experimental fMRI studies compared brain activity across two or more experimental conditions in a single group of participants. The approximate topics of experimental fMRI papers are shown in **Supplementary Table 2**.

(2) Cognitive neuroscience sMRI studies typically used structural data to support the interpretation of fMRI data; to gain anatomical information relevant for understanding normal brain function (e.g. by studying connections between areas thought to implement certain functions and/or cortical thickness in some areas thought to host some functions); to compare brain anatomy in non-clinical groups of participants (e.g. normal and poor adult readers); and to study network properties thought to support some functions.

(3-4) Clinical fMRI (3) and Clinical sMRI (4) studies with patient groups including studies of ageing and studies focused on developmental disorders in children. Many clinical MRI studies compared brain function or structure across controls and patients or measured the effect of aging by studying multiple age groups. Single group studies also tested groups of participants in various experimental conditions. In a few papers participants were healthy ‘control participants’ but the primary objective of papers was clinical research (e.g. testing the effectiveness of pain suppression). Such papers were categorized as ‘clinical’ papers. The most frequently studied diseases and conditions and associated median and mean participant numbers in clinically oriented papers are shown in **Supplementary Table 3.**

(5-6) Normative developmental fMRI (5) and sMRI (6) studies with typically developing children who were under the age of 18 years. Many of these studies compared brain function or structure across multiple age groups.

There were very few cognitive neuroscience sMRI (16 papers with 18 studies) and developmental sMRI (19 papers with 19 studies) and fMRI (25 papers with 25 studies) studies as compared with studies in the other 3 categories. Hence, data from these 60 papers (62 studies) were not considered for analysis. However, the extracted data is available as supplementary material.

Data for the remaining 1038 papers with 1161 studies were analyzed in the work reported here. These studies were categorized as experimental fMRI, clinical sMRI and clinical fMRI studies. **Table 1** shows the number of highly cited papers, the studies included in the papers and paper citation counts. Papers in this sample were published between 1990-2012 (Experimental fMRI studies: 1993-2012; Clinical sMRI studies: 1990-2011; Clinical fMRI studies: 1996-2012). The 1038 papers received 391,180 citations. The experimental fMRI papers received 231,071 of these citations.

**Table 1.** The numbers of highly cited papers, the studies included in the papers and paper citation counts. The 1038 papers were subdivided into three categories (see details below).

| Study Type | Papers | Studies | Citation Counts |  |  |  |  |
| --- | --- | --- | --- | --- | --- | --- | --- |
|  |  |  | Min | Median | Mean | Max | Total |
| **All** | 1038 | 1161 | 208 | 297 | 377 | 4147 | 391,178 |
| **Experimental fMRI** | 591 | 692 | 208 | 305 | 391 | 4147 | 231,071 |
| **Clinical sMRI** | 318 | 334 | 208 | 287 | 358 | 2912 | 113,954 |
| **Clinical fMRI** | 129 | 135 | 209 | 300 | 358 | 981 | 46,153 |

We extracted the following data for each study: 1) Total number of participants tested. 2) The number of participants stated as excluded from analyses. When no exclusions were reported we assumed that the number of excluded participants was zero. 3) The final number of participants included in the MRI analysis. This ‘final’ participant number was considered the number of participants for a study. 4) We determined whether a study defined two or more groups of participants. If at least two groups were defined then we recorded the number of participants in each group. 5) For experimental studies we noted the approximate main topic of a paper. 6) For clinical studies we coded the type of disease examined. 7) Finally, we coded whether a study was a randomized control trial or not.

### Analysis of trial numbers in highly cited experimental fMRI papers

In order to get an impression of total and per condition experimental trial numbers in individual participants we have examined the Methods sections of 142 experimental fMRI studies with event-related designs where trial numbers should be well-defined in principle. We extracted the total number of trials and the number of experimental conditions where this was possible.

### Sample of experimental fMRI papers in 2017 and 2018

We analyzed a sample of 131 experimental fMRI papers published during 2017 and 142 papers published during 2018 in 4 prominent neuro-imaging journals: *Nature Neuroscience, The Journal of Neuroscience, NeuroImage and Cerebral Cortex*. The number of studies and papers are shown above in **Table 2**. The issues checked per journal are shown in **Supplementary Table 4**.

**Table 2.** The number of papers and studies in the 2017 and 2018 sample.

| Journal | 2017 |  | 2018 |  | Totals |  |
| --- | --- | --- | --- | --- | --- | --- |
|  | Papers | Studies | Papers | Studies | Papers | Studies |
| **Nature Neuroscience** | 4 | 5 | 2 | 2 | 7 | 7 |
| **The Journal of Neuroscience** | 42 | 47 | 33 | 38 | 80 | 85 |
| **NeuroImage** | 46 | 51 | 68 | 74 | 119 | 125 |
| **Cerebral Cortex** | 39 | 45 | 39 | 40 | 84 | 85 |
| **Totals** | 131 | 148 | 142 | 154 | 273 | 302 |

During data collection we manually opened each pdf file in Adobe Acrobat © and checked the ‘Participants’ or equivalent section for initial and final sample sizes and for the number of excluded participants. In addition, we have also searched papers for the words ‘power’ and ‘sample size’ and determined whether papers included any formal power calculations and/or they justified their sample sizes. In order to verify power analyses in papers and/or to confirm their nature we have carried out our own power analyses for each paper based on the data given in the papers. This procedure was most often necessary because from papers it was not clear what kind of power analysis was exactly done.

### Data availability

All data and the analysis code (Matlab scripts; www.mathworks.com) producing all figures, tables and numerical details reported here will be available as **Supplementary Material** of the final publication. It is not possible to upload pdf copies of published papers because of copyright restrictions.

## Results

### Highly cited paper sample: 1990-2012

**Figure 1A** compares sample size distributions for the 3 types of papers in the highly cited sample. **Table 3** shows the number of participants in studies with a single group and with more than one group. For example, 662 out of 692 experimental studies had a single group of participants whereas 30 studies defined two or more groups. The median number of participants in studies with a single group was 12. In the 30 studies with groups the average group number was 2.033. The median number of participants in groups was 11. The proportion of studies with a single group is notably higher in experimental fMRI studies (662/692=0.9566) than in Clinical sMRI (163/334=0.4880) and Clinical fMRI studies (28/135=0.2074). Median participant numbers were 3.5 to 4.17 times larger (N=50) in single group Clinical sMRI studies than in the other two study categories. Median participant numbers were about twice as large in multi-group Clinical sMRI studies (N=24) than in other study categories.

**Table 3.** The number of participants in highly cited studies with a single group (gr=1) and with more than one group (gr>1). Study categories: Experimental (Exp) fMRI and Clinical (Clin) sMRI and fMRI studies. The first 3 columns show the number of studies with one or more groups and totals. The next 3 columns (N in Group if gr=1) show participant numbers for studies with a single group. The next 3 columns (Number of Groups if gr>1) show the number of groups in studies with more than one group. The last 3 columns (N in Groups if gr>1) show participant numbers in groups in studies with more than 1 group.

| Study Type | Studies |  |  | N in Group if gr=1 |  |  | Number of Groups if gr>1 |  |  | N in Groups if gr>1 |  |  |
| --- | --- | --- | --- | --- | --- | --- | --- | --- | --- | --- | --- | --- |
|  | gr=1 | gr>1 | Total | Min | Median | Max | Min | Mean | Max | Min | Median | Max |
| All | 853 | 308 | 1161 | 2 | 13 | 3660 | 2 | 2.35 | 8 | 1 | 17.2 | 1056 |
| Exp fMRI | 662 | 30 | 692 | 2 | 12 | 603 | 2 | 2.03 | 3 | 1 | 11.0 | 500 |
| Clin sMRI | 163 | 171 | 334 | 2 | 50 | 3660 | 2 | 2.50 | 8 | 2 | 24.0 | 1056 |
| Clin fMRI | 28 | 107 | 135 | 3 | 14.5 | 146 | 2 | 2.19 | 7 | 1 | 12.5 | 68 |

**Figure 1.**
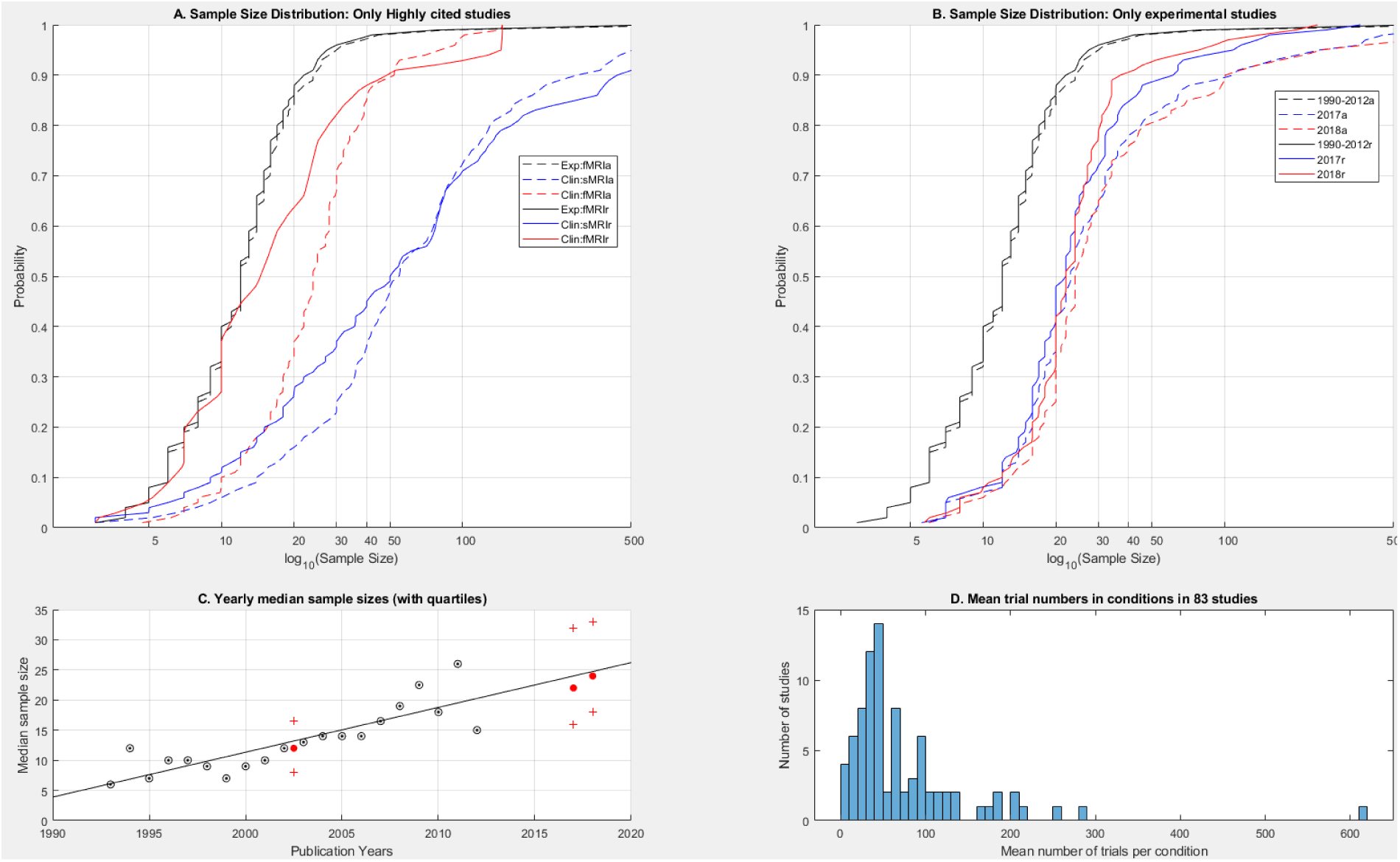
**(A)** Sample size distributions in highly cited papers. Dashed lines marked by (a) in the legend show data for all studies. Continuous lines marked by (r) in the legend show data restricted to studies with that collected their own data and only included a single group of participants. **(B)** Sample size distributions in experimental fMRI papers in highly cited papers and in 2017 and 2018. Dashed (a) and continuous lines (r) show data as noted for Panel A. **(C)** Black circled dots show the yearly medians of the sample sizes from the highly cited papers. The black line is the regression line fitted to this data. The leftmost red dot and red crosses represent the median and 25^th^ and 75^th^ percentiles of sample sizes from the entirety of the highly cited paper data. The rightmost two red dots and crosses represent the medians and 25^th^ and 75^th^ percentiles of 2017 and 2018 data. **(D)** The distribution of mean number of trials in the experimental conditions of 83 highly cited experimental fMRI papers (see further explanation in text).

There was no relationship between the number of citations to a paper and the number of participants in studies. In the whole sample of 1161 studies the correlation of citation count and sample size was r=0.0098 [95% CI: −0.0413; 0.0609; p=0.71]. In the sample of Experimental fMRI papers the correlation was r=0.0342 [95% CI: −0.0405; 0.1084; p=0.37].

### Studies from 2017 and 2018

**Table 4** shows the counts and proportion of papers and studies with their own data, with secondary data and with only one group in the 2017 and 2018 data sets. Similarly to the highly cited paper data the overwhelming majority of papers had a single group of participants.

**Table 4.**
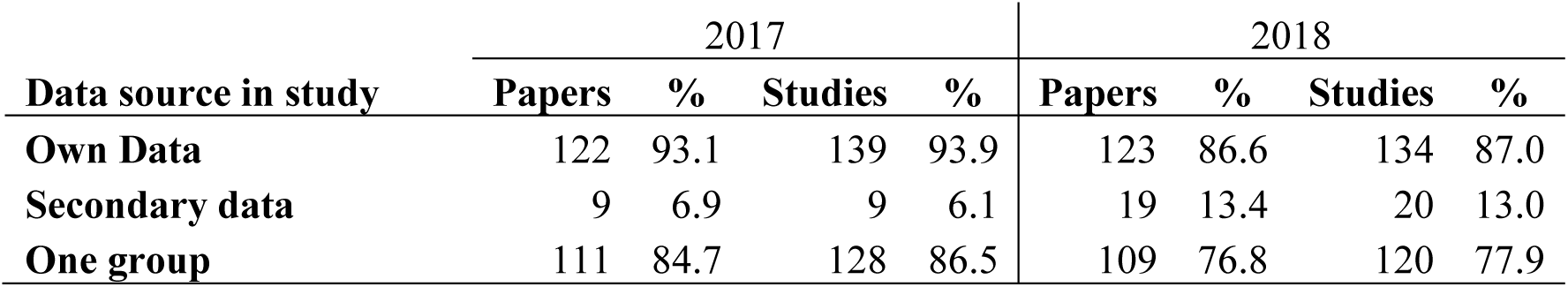
The counts and percentage of papers and studies with their own data, with secondary data and with only one group in the 2017 and 2018 data sets. The studies with one group are a subset of the studies with their own data. Percentages are computed relative to total paper and study numbers as shown in Table 2.

| Data source in study | 2017 |  |  |  | 2018 |  |  |  |
| --- | --- | --- | --- | --- | --- | --- | --- | --- |
|  | Papers | % | Studies | % | Papers | % | Studies | % |
| Own Data | 122 | 93.1 | 139 | 93.9 | 123 | 86.6 | 134 | 87.0 |
| Secondary data | 9 | 6.9 | 9 | 6.1 | 19 | 13.4 | 20 | 13.0 |
| One group | 111 | 84.7 | 128 | 86.5 | 109 | 76.8 | 120 | 77.9 |

### Comparison of sample size percentiles from highly cited papers and from papers published in 2017 and 2018

**Figure 1B** compares sample size distributions in the highly cited paper sample and in the 2017-2018 sample. The shift in sample size distributions and a slight increase in the proportion of studies with large sample sizes (most of them studies based on data from large third party data bases) is well visible (sample size percentiles are shown in **Supplementary Table 5**). **Table 5** shows what number and proportion of studies with their own data and a single group of participants exceeded certain participant numbers recommended by Desmond and Glover (2002) and Yarkoni (2009).

**Table 5.** The number and proportion (Prop.) of studies with their own data and a single group of participants that exceeded participant numbers of 12, 24, 50 and 100 (N=).

|  |  | N=12 | N=24 | N=50 | N=100 |
| --- | --- | --- | --- | --- | --- |
| **Exp fMRI** | **p>N** | 372 | 62 | 10 | 4 |
|  | **Prop. (all=662)** | 0.562 | 0.094 | 0.02 | 0.006 |
| **Clin sMRI** | **p>N** | 141 | 113 | 84 | 48 |
|  | **Prop. (all=163)** | 0.865 | 0.693 | 0.52 | 0.2945 |
| **Clin fMRI** | **p>N** | 16 | 8 | 3 | 2 |
|  | **Prop. (all=28)** | 0.571 | 0.286 | 0.11 | 0.0714 |
| **2017** | **p>N** | 117 | 53 | 14 | 7 |
|  | **Prop. (all=128)** | 0.914 | 0.414 | 0.11 | 0.0547 |
| **2018** | **p>N** | 109 | 57 | 9 | 4 |
|  | **Prop. (all=120)** | 0.908 | 0.475 | 0.08 | 0.0333 |

**Figure 1C** shows the increase in median sample sizes from 1993 to 2018. The figure shows the regression line computed from the highly cited paper data (black line). According to data form highly cited papers the rate of increase in sample sizes was +0.74 participants/year (intercept = −1477). The figure also shows the medians and 25^th^ and 75^th^ percentiles of the 2017 and 2018 data. It is notable that extrapolation of the regression line extremely well fit the medians measured in 2017 and 2018. In **Figure 1C** it is visible that medians show larger scattering relative to the regression line at the left and right extremes of data points. This is due to the fact that less data points were available in very early and very recent publication years (see numbers in **Supplementary Table 6**.)

### The number of reported excluded participants in highly cited experimental fMRI papers

**Table 6** shows the number of *reported* excluded participants per study category. 86-90% of highly cited studies did not report excluded participants. In contrast, about 45% of studies in 2017 and 2018 reported some excluded participants. The proportion of studies with a relatively large number of excluded participants (6-10 or more excluded participants) notably increased from 1-2% in highly cited studies to about 9-10% by 2017 and 2018. More excluded participants were reported in clinical than in experimental studies.

**Table 6.** The number of excluded participants per study category. Numbers are given for studies (not papers). For each category the following data are shown: total number of studies (Total), studies with no reported exclusions (None), studies with some exclusions (Some), studies with certain numbers of participants excluded (1-5, 6-10, >10). The median and mean of the number of excluded participants. For each category top rows (N) communicate participant numbers and bottom rows (%) communicate the percent of participants relative to the total participant number in a category.

|  |  | Total | None | Some | 1-5 | 6-10 | >10 | Median | Mean |
| --- | --- | --- | --- | --- | --- | --- | --- | --- | --- |
| **Exp**<br>**fMRI** | N | 692 | 595 | 97 | 80 | 13 | 4 | 2 | 3.7 |
|  | % | 100 | 86 | 14 | 12 | 2 | 1 |  |  |
| **Clin**<br>**sMRI** | N | 334 | 301 | 33 | 6 | 7 | 20 | 22 | 87.5 |
|  | % | 100 | 90 | 10 | 2 | 2 | 6 |  |  |
| **Clin**<br>**fMRI** | N | 135 | 121 | 14 | 8 | 3 | 3 | 4 | 8.4 |
|  | % | 100 | 90 | 10 | 6 | 2 | 2 |  |  |
| **2017** | N | 148 | 80 | 68 | 47 | 13 | 8 | 3 | 5.0 |
|  | % | 100 | 54 | 46 | 32 | 9 | 5 |  |  |
| **2018** | N | 154 | 84 | 70 | 41 | 15 | 14 | 4 | 15.1 |
|  | % | 100 | 55 | 45 | 27 | 10 | 9 |  |  |

### Trial numbers in a sample of highly cited experimental papers

We could identify total and per condition trial numbers in 109 of the 142 experimental fMRI papers, while this information was unclear in the other 33 papers.

17 papers described old/new recognition memory experiments. We extracted the number of memory encoding trials as the usual question of interest is whether some brain activity at encoding will predict later recognition. The number of trials ranged from 10 to 455 (median = 150). 9 papers described designs with a large number of standard trials interspersed with a significantly lesser number of deviant trials from a different, critical trial type, for example in go/nogo designs (where typically there are many fewer nogo than go trials) and in task switch designs (where typically there are much fewer task switch than standard trials). Trial numbers varied from 128 (8 critical) trials to 1180 (80 critical) trials.

In 83 of the 109 papers trial numbers were more similar across conditions than in the above standard/deviant like designs. However, trial numbers were still very often unequal across conditions and there was also great variability in design. **Figure 1D** shows the mean number of trials by condition. The number of total number of trials in an experiment ranged between 40 and 2440, the number of conditions ranged between 2 and 28 and the mean number of trials per condition ranged between 4 and 610. For example, on the one extreme 112 trials were distributed into 28 conditions and on the other end 2440 trials were distributed into 4 conditions. It is notable in the figure that the mean number of trials per condition tends to decrease as the number of experimental conditions increases.

### Power calculations in 2017 and 2018 papers

**Table 7** shows the summary of the statistical power analysis assessment. In both 2017 and 2018 less than 7% of papers (9 papers in both years) included power calculations and about a third of the papers made some comment about power. None of the papers with large secondary databases had power calculations in any of the years. 7.6% (10) vs. 11.3% (16) of papers without power calculations referred specifically to the problem of having low power, in 2017 and 2018, respectively. Only about 3% of 2017 and 2018 papers had clearly a priori power calculations.

**Table 7.** Summary of power calculation results. The table shows data for the 131 papers in 2017 and the 142 papers in 2018. It is shown whether papers included statistical power calculations (a), had any comments on power (b) or had no comments on power (c). It is also shown whether power computations were a priori or post-hoc if we could determine this.

| Information on power? | 2017 |  | 2018 |  |
| --- | --- | --- | --- | --- |
|  | N | % of 131 | N | % of 142 |
| **Power calculation (a)** | 9 | 6.9 | 9 | 6.3 |
| - A priori power calculation | 4 | 3.1 | 6 | 4.2 |
| - Post-hoc power calculation | 3 | 2.3 | 3 | 2.1 |
| - Unclear | 2 | 1.5 | -- | -- |
| **Comments on power (b)** | 34 | 26.0 | 44 | 31.0 |
| **No comments on power (c)** | 88 | 67.2 | 89 | 62.7 |

### Power calculation details in 2017 and 2018 papers

In 2017 four papers seemed to include formal pre-study power calculations (α=0.05 for all). Two of these papers computed power for single runs of t-tests (two cases; Cohen’D =0.5 and 0.65; power=0.8 for both). Two other distinct papers (n=32) in the same journal issue used the same participants. One paper computed power for a single one-sample t-test (D=0.5) and set a priori power to exceed 0.85. The other computed power for a single product-moment correlation (r=0.4) and set a priori power to exceed 0.75.

In 2018 six papers described a-priori power computations. *One paper* determined that a sample size of 24 was necessary to achieve power=0.8 with a matched-sample two-tailed t-test to detect an effect size of D ≥ 0.6 (α=0.05). However, 2 participants were excluded from analyses leaving only 22 participants in the study. This obviously left the study underpowered by its own power criterion. *Another paper* aimed to look at brain structure vs. behavioral performance correlations. It was not specified how exactly power computation was done but our own power analysis suggested that the required sample size of 34 was computed for power=0.8 for a single correlation test of r=0.4 (α=0.05; one-tailed). The study noted that power was computed for D=0.4 in G-Power. However, G-Power computes the sample size of 24 when r=0.4 rather than when D=0.4 (entering r is the default in G-Power). For D = 0.4; r would be r = D / sqrt(D^2^ + 4) = 0.1961 (Borenstein et al. 2009). To detect an effect size of r=0.1961 G-Power computes that N=156 participants would be necessary (α=0.05; one-tailed). For this effect size (r=0.1961) the study achieved power=0.24 with N=24 (α=0.05; one-tailed). Hence, the study had much less power than reported. *Another paper* stated that they chose a sample size to achieve good power to detect the typical effect size in the field. An effect size of r = 0.54 was chosen from a previous meta-analysis. It was concluded that for the 25 participants initially tested power=0.87 would be achieved for the D=0.54 effect size (α=0.05; two-tailed; power was computed for a single correlation). The paper initially tested 25 participants but one participant was excluded, so only 24 participants were tested.

*Another 2018 paper* computed that 34 participants were necessary to detect an interaction effect size of partial eta^2^ = 0.06 at power=0.8 and 12 participants were required to replicate a previously found effect size of partial eta^2^ = 0.17 at power=0.8. The study tested 34 participants. Based on recomputing power in G-Power it seems that power was computed for a 2(groups) x 2(measurements) mixed design ANOVA with partial eta^2^ of 0.06 and 0.17 (transformed to Cohen’s f=0.2526 and f=0.4225 by G-Power). *Another paper* referred to a previous study of the authors where they used Monte-Carlo simulation to estimate sample size. The current study aimed to test a sample size of 30 participants as required by this previous power calculation. Based on the calculation the authors concluded that they achieved power > 0.95. However, 2 participants were excluded, so only 28 participants were tested. *One paper* estimated the required sample size for Multi Voxel Pattern Analysis by a simulation method.

In 2017 in two cases it was unclear whether power was computed a priori. In these papers power (set to 0.8) was computed for a t-test (D=0.44) and for correlation (r=0.5). The first of these papers presented the only RCT in our 2017 sample with 128 participants.

In 2017 in three cases power was computed post-hoc. In these papers power was computed for a t-test (D=0.6), for an ANOVA interaction term (D=0.52) and in one case it was unclear how power was computed as no exact effect sizes were given (but likely for multiple t-tests). One of these studies (n=2×15) computed power to guide future studies.

In 2018 power computations seemed clearly post-hoc in three cases. *One paper* with 15 participants has achieved null results in whole brain analyses. The study has computed the achieved power for multiple testing uncorrected ROI analyses suggesting that analyses achieved power=0.8 to detect an effect size of D=0.68 with α=0.05 and D=0.56 at α=0.1. Computations were not specified in the paper but re-analysis suggests that they were done for one-tailed t-tests. *The second paper* noted that the study was ‘adequately powered to detect large effects’ and noted that the sample size of 12 was adequate to detect an effect size of d ≥ 0.89 at power=0.8 (α=0.05). Power was computed for a single two-tailed matched-sample t-test. *In the third paper* it was unclear how power was computed but the authors claimed to have run some analyses for effect sizes of D=0.51, 0.53 and 0.54 with power=0.8. The analyses close to describing the power computation mentioned the use of matched sample t-tests and the study had 31 participants. Indeed, power for a two-tailed t-test (α=0.05) for the above effect sizes and sample size varies between power=0.78 to 0.83. So, it is likely that power was computed for single matched-sample t-tests.

Besides the papers with power calculations 34 and 44 papers mentioned statistical power in 2017 and 2018, respectively. Mentions were most often non-specific and non-informative. For example, all but one paper in Nature Neuroscience had power calculations but all of them included similar text stating that no methods have been used to predetermine sample sizes and sample sizes were simply chosen to be in line with common practices in their field. Many studies noted that their sample size was chosen so that they would be identical to, or exceed sample sizes from the authors’ own or others’ previous work. Some noted that sample size was based on funding availability. Some studies commented on issues of statistical power in general without clearly linking it to the context of the specific study. Some studies commented that a certain analysis was less or more powered than another one without giving any further details or computations. Some papers commented on their large perceived sample size (e.g. 60) without giving any actual power calculation details. Ten (2017) and sixteen (2018) studies mentioned that they may have been underpowered. Usually these non-informative statements were restricted to one or two brief comments in a paper. Several studies used small subsamples from their overall sample for certain analyses.

In 2017 seven whereas in 2018 only two papers had multiple studies where some of these studies declared a goal to replicate findings from an earlier study in the same paper. None of these papers had power calculations.

In 2017 two studies were special cases focused on individual measurement: One had only 4 participants but each of them were tested in 5-6 sessions. The other tested 10 participants, each for 300 minutes during 10 sessions.

## Discussion

Running underpowered studies may waste research funding on studies which a priori have low chance to achieve their objectives. In addition, low power leads to high false report probability, imprecise measurements and effect size exaggeration. Nevertheless, we have shown that participant numbers, and consequently power levels, are low in the most highly cited fMRI papers. Hence, highly cited studies are likely to have similar problems stemming from low statistical power as most of ‘typical’ neuroscience studies (Button et al. 2013; Szucs and Ioannidis, 2017a).

Highly cited experimental and clinical fMRI studies had similar median sample sizes (medians in single group studies: 12 and 14.5; median group sizes in multiple group studies: 11 and 12.5). 96% of experimental studies were single group studies. This pattern remained in 2017 and 2018 when 93% and 87% of experimental fMRI studies had a single group. In contrast, only 21% of highly cited clinical fMRI studies were relatively small single group studies. Consequently, while clinical fMRI studies had somewhat larger individual participant groups than experimental studies, most of their total sample size advantage stemmed from the fact that they more often had two or more groups than experimental studies. This is important to consider when comparing sample sizes from clinically vs. non-clinically oriented journals/publications. Clinical studies included multiple groups because these studies often included both patient and control groups. Overall, group sizes were relatively similar in both experimental and clinical fMRI studies. In addition, group comparison tests are typically less powerful than one-sample or matched sample-tests often used in single group studies. Hence, in terms of group comparison there was not much power advantage of clinical fMRI studies over experimental fMRI studies. Only the single-group clinical sMRI studies had notably larger sample sizes than fMRI studies. This difference probably has to do with the extra time and effort necessary to collect and analyze fMRI than sMRI data.

We found that median sample sizes in highly cited experimental fMRI studies increased consistently at a rate of +0.74 participant/year between 1993 and 2010. This rate of increase was perfectly in line with the median sample sizes we found in 2017 (23) and 2018 (24). The +0.74 participant/year rate of increase was also in line with our survey (Szucs and Ioannidis, 2017a) examining 3801 papers published between 2011 and 2014. In this earlier paper we reported degrees of freedom for one or two-sample t-tests and estimated that the median degree of freedom was 18 in cognitive neuroscience papers. Provided that median sample sizes were likely to be about 1-2 larger than the degrees of freedom this data would also well fit the regression line found in the current study (with about median sample size of 20-21 in about 2012/13). Notably, our sample size median estimates are smaller than the 28.5 median estimated by Poldrack et al. (2017) for the year of 2015. However, our analysis is well compatible with the full set of data points from David et al. (2013) reported by Poldrack et al (2017). An option is that the 2015 data from Poldrack et al. (2017) may have included a higher proportion of clinical fMRI papers than our sample which may have raised sample sizes for 2015.

Overall, our current and earlier data (Szucs and Ioannidis, 2017a) and data from other evaluations (David et al. 2013; Poldrack et al. 2017) suggest that sample sizes and consequently, power are improving, albeit very slowly. Only ∼10% of highly cited experimental fMRI papers published between 1993-2012 reached the sample size of 24 recommended by Desmond and Glover (2002) and only about ∼2% reached the sample size of 50 recommended by Yarkoni (2009). There has been clear improvement by 2017 and 2018 when respectively, 41% and 48% of papers were below the minimum participant numbers recommended by Desmond and Glover (2002). However, this also means that in 2017 and 2018 still nearly half of papers had less than 24 participants. Moreover, there has been less improvement at the higher end of participant numbers as in 2017 still 79% of papers were under the sample size of 50 recommended by Yarkoni (2009) and this proportion was 82% in 2018.

While we only have two years’ of observations from 2017 and 2018 a noteworthy trend in the literature is the increasing use of large third party databases in neuroimaging. The proportion of papers using such databases doubled from 2017 to 2018 from 6% of studies to 13% of studies. It remains to be seen whether this trend continues in the coming years and whether it is present in other neuroimaging journals. The use of large shared databases would be beneficial for many reasons. First, such databases assure high statistical power for modest effects. Second, considering the effort required to compile large databases data collection may be carried out by seriously vetted procedures and by experienced teams. Third, data is available to any interested researchers assuring increased scrutiny and hence, reliability of published results. Fourth, if data is collected in a decentralized manner (e.g. many labs jointly contributing to data collection) than replicability across different labs can easily be examined.

While low power in neuro-imaging received lots of attention recently (Poldrack et al. 2017; Szucs and Ioannidis, 2017; Button et al. 2013), we found that in both 2017 and 2018 only about 3-4% of papers had clear pre-study power calculations and more than 62% of papers never mentioned any issues of statistical power. Most power calculations we found were done for single runs of t-tests and product-moment correlations as there seems to be no agreement on how to estimate statistical power for fMRI studies which rely on a very large number of tests, idiosyncratic statistical procedures and on heavy multiple testing correction (Hayasaka et al. 2007; Poldrack et al. 2017, Carp 2012). The power calculations we found often expected medium sized effects based on previous published data. However, considering the very probable effect size inflation of the published literature expecting relatively large medium sized effects seems too optimistic (Ioannidis, 2008; Szucs and Ioannidis, 2017a). It also frequently happened that studies determined a required sample size by power calculation but then analyzed less data than required by their own power calculation because they did not account for the number of excluded participants. This practice leaves studies underpowered by their own power criteria. It was also typical that power calculation parameters were not defined clearly so that in most cases guesswork and recalculation was necessary to see how power was determined. In some cases power calculations seemed erroneous.

Many papers without power calculations referred to sample sizes in previous similar research to justify their sample sizes. However, considering that lots of neuroimaging is underpowered (Yarkoni 2009; Button et al. 2013; Szucs et al. 2017a) this is clearly inadequate rationale. Unless the purpose is to guide future studies it is not informative to compute post-hoc power considering an effect size already detected as statistically significant in a study. Moreover, small studies are not good guides for power calculations for future studies because they can *only* detect relatively large effects as statistically significant (see e.g. Ioannidis, 2008; Yarkoni 2009; Szucs and Ioannidis, 2017b). Similarly, meta-analyses may also overestimate effects because they tend to rely on many small, underpowered studies (Ioannidis, 2010). In fact, studies with large sample size (and hence, with more accurate measurements than small studies) rarely report large effects (see Fig. 2. in Szucs and Ioannidis 2017a).

We suggest that it would be desirable to quickly develop ‘industry standards’ for technical aspects for neuroimaging studies including power requirements. In our opinion two key ingredients of future studies are pre-registration (optimally, with pre-study acceptance by journals) and an increase in sample sizes (Hardwicke and Ioannidis, 2018; Munafo et al. 2017; Ioannidis et al. 2014). Pre-registration guarantees that studies get published based on pre-study significance. In consequence, we avoid effect size exaggeration (as negative findings are published) and registered procedures and hypotheses largely decrease the danger of data dredging which is a particular danger in neuro-imaging where often complicated and opaque procedures are used and seeming minor changes in some (pre-)analysis parameters can result in major distortions of data (Carp 2012).

If expected effect sizes are larger in clinical than in experimental studies (as disease is likely to have more substantial impact on brain function than experimental manipulations in healthy participants), the power advantage of clinical over experimental studies may be larger than indicated here. Importantly, both here and earlier (Szucs and Ioannidis, 2017a) we detected considerable variability in sample sizes across studies. So, using solely medians to characterize sample sizes seems inadequate as it masks variability. The crucial question is what proportion of studies in the literature remain too small and hence, underpowered.

A further point to discuss regards the question of whether high population level power is always necessary for studies. We agree that small N designs may be more appropriate than large samples in some situations (Smith and Little, 2018). For example, if we study some relatively straightforward to localize motor or perceptual processes with already well-known brain anatomy and probably modest individual anatomical variability then using very high trial numbers in a few participants and fitting models to high volumes of data may be a more productive approach than testing large populations with a few trials. However, such design may be less adequate when researchers aim to study harder to localize processes with probably large individual variability such as autobiographical memories, emotional evaluation of stimuli, love, etc. In any case, small N designs require us to assure very high individual level statistical power as well as to demonstrate the replicability of each experimental effect at the individual level. Considering that signal to noise ratio is proportional to the square root of trials used in experiments assuring the above would require us to collect much higher volumes of individual data than typical at the moment. This could be achieved by delivering a very high number of trials per experimental condition and participant substantially prolonging experimental time for individual participants. The challenge is made even more difficult by natural variation in the anatomical location of effects. However, as shown in Results, currently very few studies deliver high trial numbers. Hence, currently very few studies can argue that they use credible small N designs: Simply having small N does not turn a study into a credible small N study. In fact, probably due to low individual level signal-to-nose ratio it is a well-known problem of neuroscience experiments that it is often difficult to identify even well-established group level effects in individual participants. Further, even if we use large trial numbers but only a very few participants then individual measurements will be very precise but we may still not be able to generalize anatomical findings (that are often the focus of fMRI studies) to the full population. For example, if we measure a phenomenon with small individual standard errors in 4 undergraduate university students (e.g. using 5,000 trials per condition in each participant) but individual brain activity measures and anatomical areas show large scattering then we cannot approximate the population distribution of anatomical brain activation locations with much confidence. Even if individual measurements are very similar in a few participants, functional and anatomical generalizability remains unknown. For example, non-clinical experimental results are very often coming from “WEIRD” undergraduate students from Western, Educated, Industrialized, Rich and Democratic countries. A potential approach for increasing confidence in group level findings may be to replicate a finding from a key experiment in a (pre-registered) follow-up experiment within the same paper. However, this practice is currently also very rare (see Results) and in some cases it may also be difficult to assure the independence of replication experiments reported within a single paper.

Importantly, the number of trials in single fMRI measurement sessions is constrained by various practical factors. First, the sluggish nature of the haemodynamic response requires relatively long trial durations (e.g. much longer than in electro-encephalography experiments). Second, fMRI scanning time is expensive (e.g. costs may be higher than 500/hour in the United Kingdom). Third, lying in the scanner relatively motionless is challenging for participants. Fourth, complex cognitive experiments may need long trial durations which restrict the number of trials doable in a single imaging session. The only solution may be to collect many trials in multiple runs. However, in such a case multi-level analysis is needed to factor in potential discrepancies across sessions. Moreover, research grants can rarely offer funds for long/repeated scanning sessions. Hence, overall several practical limitations often beyond the control of researchers restrict individual measurement precision in studies. Finally, our data shows that individual research groups may use very different trial numbers even in relatively similar experimental designs and overall there is very great variability in trial numbers per experimental condition in the literature. Hence, individual measurement precision is likely to vary greatly across studies and research groups.

Only 10-15% of highly cited studies reported any excluded participants. In contrast, in 2017 and 2018 respectively, 46% and 45% of studies reported at least some excluded participants. The proportion of studies with 5 or more excluded participants also increased by about 5-fold by 2017/2018. Importantly, studies practically never stated that *no* participants were excluded. Hence, the default value was that studies have not mentioned anything about exclusions. That is, when there were no reported exclusions it may mean that there were really no exclusions or that exclusions were simply not reported. Taking the above into account our observations about reporting exclusions raise several questions. On the one hand, the distribution of excluded participants in highly cited papers seems oddly biased towards 0 exclusions and it is also in conflict with the much larger proportion of studies with exclusions and the larger number of excluded participants in 2017/18. So, many highly cited studies may not have reported exclusions rather than not have exclusions. If so, there may have been a change in exclusion reporting habits during the past years, Alternatively, perhaps the most recent papers do exclude more participants then earlier papers. Overall, it is not possible to decide which of the above scenarios may be more probable. However, it is important to note that exclusions do allow for high ‘researcher degree of freedom’ and so have implications for data dredging and N-hacking (Simmons et al. 2011; Carp 2012; Szucs 2016). Therefore, it would be important to clarify and define exclusion criteria and numbers in research fields.

Besides the uncertainty about interpreting the above exclusion data it is also noteworthy that extracting condition numbers and trial numbers in each condition in published papers was particularly difficult due to completely idiosyncratic descriptions and missing information. So, we suggest that it would be beneficial to standardize reporting requirements and formats in neuroimaging, for example by linking standard reporting cards to *all* neuroimaging papers. For example Nature Research has already started using standard ‘Reporting Summaries for MRI studies’. However, we suggest a more formal, comprehensive and universally required approach (see related discussion in Begley and Ioannidis, 2015). Standardized reporting would also make papers easily machine readable, with their data being possible to re-analyze and to combine.

In conclusion, the consistent historic increase in sample sizes suggests that we may be able to break the long ‘tradition’ of criticizing low power but not improving the situation (Sedlmeyer and Gigerenzer, 1989). However, the increase in sample sizes could be sped up by targeted and timely interventions by both publishers and funders. Funding contracts could specify power-calculation-based sample sizes, pre-registration requirements for crucial studies and standardized reporting of methods/results. Such changes would provide funders with certainty that their money is not wasted.

It is tempting to assume that many of the highly cited papers analyzed here are probably replicated given that so many other scientists cite them. However, high citations are not synonymous to replication. It is well known from other fields that some papers get extremely heavily cited without any attempt to replicate them and that when replication eventually is attempted, it fails (Ioannidis 2007). A survey of the most-highly cited papers across all medicine has shown that of the most-cited observational studies 5 out of 6 were subsequently refuted and even a quarter of randomized trials were contradicted (Ioannidis 2005). Exact replication in particular is often avoided and this may allow building large literatures upon questionable findings. Therefore, we suggest that, whenever this has not been done already, the *exact* replication of some highly cited influential studies should be high priority as many of these ground-breaking studies were done in a previous era with deficient sample size standards. Consequently, some highly cited studies in this field may have high false report probability (Yarkoni 2009; Szucs and Ioannidis, 2017a,b).

## Acknowledgements

The study received funding from the James S McDonnell foundation through the James S. McDonnell Foundation 21^st^ Century Science Initiative in Understanding Human Cognition – Scholar Award (DS; No 220020370). METRICS is supported by a grant from the Laura and John Arnold Foundation. The work of John Ioannidis is supported by an unrestricted gift from Sue and Bob O’Donnell. We thank Timothy Myers for an initial check on sample sizes in about half the highly cited papers. We thank Josefína Weinerova for extracting sample size data for 2018. We thank Rik Henson (University of Cambridge) for comments on an earlier version of this manuscript.

